# Use of Phenylacetonitrile Plus Acetic Acid to Monitor *Pandemis pyrusana* (Lepidoptera: Tortricidae) in Apple

**DOI:** 10.1101/092452

**Authors:** A. L. Knight, E. Basoalto, G. J. R. Judd, R. Hilton, D. M. Suckling, A. M. El-Sayed

## Abstract

A recent discovery have demonstrated that herbivore induced plant volatile compounds from apple tree infested with leafrollers were highly attractive to con-specific adult male and female leafrollers. However, this work has been conducted in New Zealand and Canada testing only low doses of kairomone. This study has been conducted in US to assess the attractiveness of higher doses of the six apple volatiles provisory identified in apple trees infested by tortricid larvaeto the leafroller, *Pandemis pyrusana* Kearfott. These volatiles included, β-caryophyllene, germacrene D, benzyl alcohol, phenylacetonitrile, (*E*)-nerolidol, and indole. No volatiles were attractive to *P. pyrusana* when used alone. However, traps baited with phenylacetonitrile plus acetic acid caught both sexes of *P. pyrusana*. Traps baited with the other volatiles plus acetic acid caught zero to only incidental numbers of moths, ≤ 1.0. Adding phenylacetonitrile to traps baited with pear ester, ethyl (*E,Z*)-2,4-decadienoate plus acetic acid significantly reduced catches of codling moth, *Cydia pomonella* (L.). However, adding phenylacetonitrile to traps baited with codling moth sex pheromone, pear ester, and acetic acid did not similarly reduce moth catches of *C. pomonella*. Interestingly, traps baited with phenylacetonitrile plus acetic acid caught significantly more *P. pyrusana* than traps baited with a commercial sex pheromone lure. The evaporation rate of the acetic acid co-lure was an important factor affecting catches of *P. pyrusana* with phenylacetonitrile, and studies are needed to optimize the emission rates of both lure components. Further studies are warranted to develop phenylacetonitrile and possibly other aromatic plant volatiles as bisexual lures for the range of tortricid pests attacking horticultural crops.

---

*Pandemis* spp. leafrollers are an important group of tortricid pests attacking pome fruits (apple, *Malus domestica* Borkhausen, and pear, *Pyrus communis* L.), in North America (Chapman 1973, Dombroskie and Sperling 2012) and in Europe and Asia (Dickler 1991). These species are often bivoltine and overwintering larvae in the fall-spring and summer generations feed on the skin of developing fruits adjacent to leaves creating cull fruits (Brunner 1996). Insecticides are most often used to manage these pests, but sex pheromone mating disruption has been developed and is implemented on a relatively small acreage in North America (Knight and Turner 1999, Judd and Gardiner 2008). Sex pheromone-baited traps are used to monitor adult populations to assess both density and phenology, but pest managers feel they are insensitive to local population densities within the orchards (Brunner 1984). Alternative lures, including acetic acid have been evaluated to monitor both sexes of *Pandemis pyrusana* Kearfott (Knight, 2001); however, acetic acid alone was considered to be a weak lure and was not adopted by Washington State growers to monitor *P. pyrusana* (Alway 2003).

Several common pome fruit volatiles typically released by healthy or insect-infested fruit and/or foliage in combination with acetic acid have been evaluated as attractants for *P. pyrusana* and several other key tortricid pests (Knight et al. 2014). Volatiles tested included ethyl *(E,Z)-* 2,4-decadienoate (pear ester), (*E*)-β-farnesene, (*E*)-4,8-dimethyl-1,3,7-nonatriene, (Z)-3-hexenyl acetate, and (*E,E*)-farnesol. Only pear ester and (*E,E*)-farnesol significantly increased total moth catch (a mix of *P. pyrusana* and *Choristoneura rosaceana* (Harris)) when added to traps baited with acetic acid. Subsequently, traps baited with pear ester, (*E*)-4,8-dimethyl-1,3,7-nonatriene, butyl hexanoate, or (*E*)-β-ocimene each in binary combination with acetic acid did not catch significantly more *P. pyrusana* than traps baited with acetic acid alone (Knight et al. 2014). Nevertheless, a novel approach using traps baited with (*E,E*)-8,10-dodecadien-1-ol, the sex pheromone of codling moth, *Cydia pomonella* (L.), pear ester, and acetic acid together was evaluated as a technique to simultaneously monitor codling moth and *P. pyrusana* (Knight et al. 2014). The secondary catch of *P. pyrusana* in these traps characterized local population densities of *P. pyrusana* immature stages in commercial orchards more accurately than the industry standard sex pheromone-baited traps. Yet, in general, counts of leafroller adults caught in these traps were low which likely reduces the precision of this approach in estimating local leafroller populations across a range of pest densities. Thus, research has continued to focus on identifying more effective attractants for adult *P. pyrusana*.

Apple seedlings uniquely released several compounds including acetic acid, acetic anhydride, benzyl alcohol, benzyl nitrile, indole, 2-phenylethanol, and (E)-nerolidol only when infested by larvae of light brown apple moth (LBAM), *Epiphyas postvittana* (Walker), the obliquebanded leafroller (OBLR), *Choristoneura rosaceana* (Harris) and the eye-spotted bud moth (ESBM), *Spilonota ocellana* (Denis & Schiffermüller) (Suckling et al. 2012; El-Sayed et al. 2016). Recently El-Sayed et al. (2016) found that binary blend of the two HIPVC, benzyl nitrile + acetic acid and 2-phenylethanol + acetic acid attracted a significant number of conspecific male and female adult LBAM in New Zealand. Further investigation with other leafrollers (Tortricidae) in Canada including the ESBM and OBLR revealed similar systems. The study reported by El-Sayed et al. 2016 targeted mainly ESBM and OBLR using a lower doses of the new kairomone. In this study we have investigated the response of *P. pyrusana* to the compounds reported in El-Sayed et al. 2016 using higher doses of the HIPVC in Washington State during 2012-2014. Also, studies were conducted to evaluate possible interactions with any of the attractive volatiles when combined with pear ester and acetic acid in binary and ternary blends for moth catches of both *C. pomonella* and *P. pyrusana* in the same trap.

## Materials and Methods

### Lures

Chemical purity of the volatile apple chemicals used in our trials were as follows: benzyl alcohol (99%), (*E*)-nerolidol (85%), phenylacetonitrile (99%), indole (99%), β-caryophyllene (97%), and germacrene D (96%) (Sigma Aldrich, St. Louis, MO). These chemicals were loaded (100 mg) into plastic sachet lures. Sachet lures consisted of a heat-sealed, semi-permeable polyethylene bag (45 mm × 50 mm, 150 μm wall thickness), with a piece of cellulose acetate filter (15 mm × 45 mm, Moss Packaging Co. Ltd., Wellington, New Zealand) inserted as the carrier substrate. Two acetic acid (glacial acetic acid (99.7%), Sigma Aldrich) co-lures were made by drilling either 1.0 or 3.2 mm holes in the cap of 8 ml Nalgene vials (Nalg-Nunc International, Rochester, NY) and loading each vial with two small cotton balls and 5 ml of glacial acetic acid. A third acetic acid lure, a proprietary round (3.4 cm diameter) plastic membrane cup (Pherocon AA) was provided by Trécé Inc. (Adair, OK). The mean weekly weight loss (evaporation rate) of acetic acid from the three lures was recorded from 30 August to 26 September 2013. Individual lures (N = 10) were placed in red Pherocon VI delta traps (Trécé Inc.) without sticky liners that were hung in the canopy of five linden trees, *Tilia cordata* Miller, at the Yakima Agricultural Research Laboratory, Wapato, WA. Similarly, the weight loss of sachet lures loaded with phenylacetonitrile (N = 5) was recorded from 28 August to 19 September 2013. In addition, the proprietary lures: Pherocon CM-DA lure loaded with pear ester, the Pherocon CM-DA combo lure loaded with codling moth sex pheromone and pear ester, and the Pherocon PLR lure (#3147) loaded with the *P. pyrusana* sex pheromone were obtained from Trécé Inc.

### Field experiment protocols

Studies evaluated single, binary, and ternary blends of selected host-plant volatiles and acetic acid co-lures using a standardized protocol in 2012 and 2013. Studies were conducted in two apple orchards, the U.S.D.A. research farm situated east of Moxee, WA (46^°^30’N, 120^°^10’W), and in a commercial orchard located southwest of Naches, WA (46^°^43’N, 120^°^42’W). Orange delta traps with sticky inserts (Pherocon VI, Trécé Inc.) were used in all studies. Five replicates of each lure treatment were randomized and spaced 20 – 30 m apart. Traps were placed at a 2-m height in the canopy. All lures were placed on the liner. Acetic acid vials with 3.2-mm holes were used in all studies unless specified otherwise. Traps baited with a blank sachet lure were used as a negative control in each study. Studies were repeated a variable number of times and typically each test ran for 5 – 7 d. When studies were repeated on different dates all traps were collected, new lures were transferred to new liners, and traps were re-randomized in the field. Moths were sexed in all studies.

### Comparison of induced volatiles

Plant volatiles reportedly induced by tortricid larval feeding on apple (El-Sayed et al., 2016) was evaluated in two consecutive studies in the Moxee orchard during 2013. The first study compared moth catches from 12 – 19 August in traps baited with phenylacetonitrile, benzyl alcohol, indole, (*E*)-nerolidol, β-caryophyllene, or germacrene D; and the combination of phenylacetonitrile with an acetic acid co-lure. The second study compared these same six volatiles all in combination with acetic acid co-lures against acetic acid alone from 19 – 28 August.

### Combination lures with phenylacetonitrile

Studies conducted in both 2012 and 2013 evaluated phenylacetonitrile alone, and in combination with either pear ester or acetic acid, pear ester with acetic acid, and phenylacetonitrile plus pear ester and acetic acid. Traps in 2012 were initially placed in the Moxee orchard on 1 August and traps were checked and re-randomized on 8 August. This study was terminated on 13 August and traps were moved to the Naches orchard. Traps were checked and re-randomized in this orchard on 22 and 30 August, and the study was terminated on 12 September. This same study was repeated in 2013 from 20 – 27 June at both the Moxee and Naches orchards and from 5 - 12 and 17 – 27 August in the Moxee orchard. A third study was conducted from 7 – 21 July 2014 in the Moxee orchard to evaluate the use of phenylacetonitrile with the sex pheromone / pear ester combinational lure and the acetic acid cup lure. A final study was conducted in the Naches orchard from 30 August to 10 September 2014, to compare the magnitude of the total bisexual moth catches of *P. pyrusana* in traps with phenylacetonitrile plus acetic acid with male catches in traps baited with *P. pyrusana* sex pheromone.

The importance of the acetic acid evaporation rate was evaluated over three time periods in the Moxee orchard during 2013. Traps baited with phenylacetonitrile sachet lures plus acetic acid vials with either 1.0- or 3.2-mm holes or the Pherocon AA cup lure were compared. New lures were used in each of three separate studies which were conducted from 28 August – 4 September, 4 – 11 September, and 12 – 19 September, 2013.

### Statistical analyses

Count data were subjected to a square root transformation prior to analysis of variance (ANOVA) to normalize the variances. A randomized complete block design was used in most studies with date as the blocking variable. Following a significant *F*-test in the ANOVA, means were separated with Tukey’s HSD test with an alpha value of 0.05. Treatments with zero catches or only incidental catch were not included in these analyses.

## Results

### Comparison of induced volatiles

No *P. pyrusana* were caught with any of the six plant volatiles during test 1 in the Moxee orchard (Table 1). In comparison, small numbers of moths were caught when acetic acid was added as a co-lure to traps baited with phenylacetonitrile (Table 1). During the second test comparing these six volatiles in combination with acetic acid no moths were caught with β-caryophyllene, germacrene D, or acetic acid alone. Incidental moth catches occurred in blank traps and in traps baited with (*E*)-nerolidol lures. Means > 1 moth per trap were caught with acetic acid plus either benzyl alcohol, indole or phenylacetonitrile. The bisexual total and female moth catches were significantly greater with acetic acid plus phenylacetonitrile than the other two volatiles plus acetic acid co-lure combinations (Table 1).

**Table 1.**
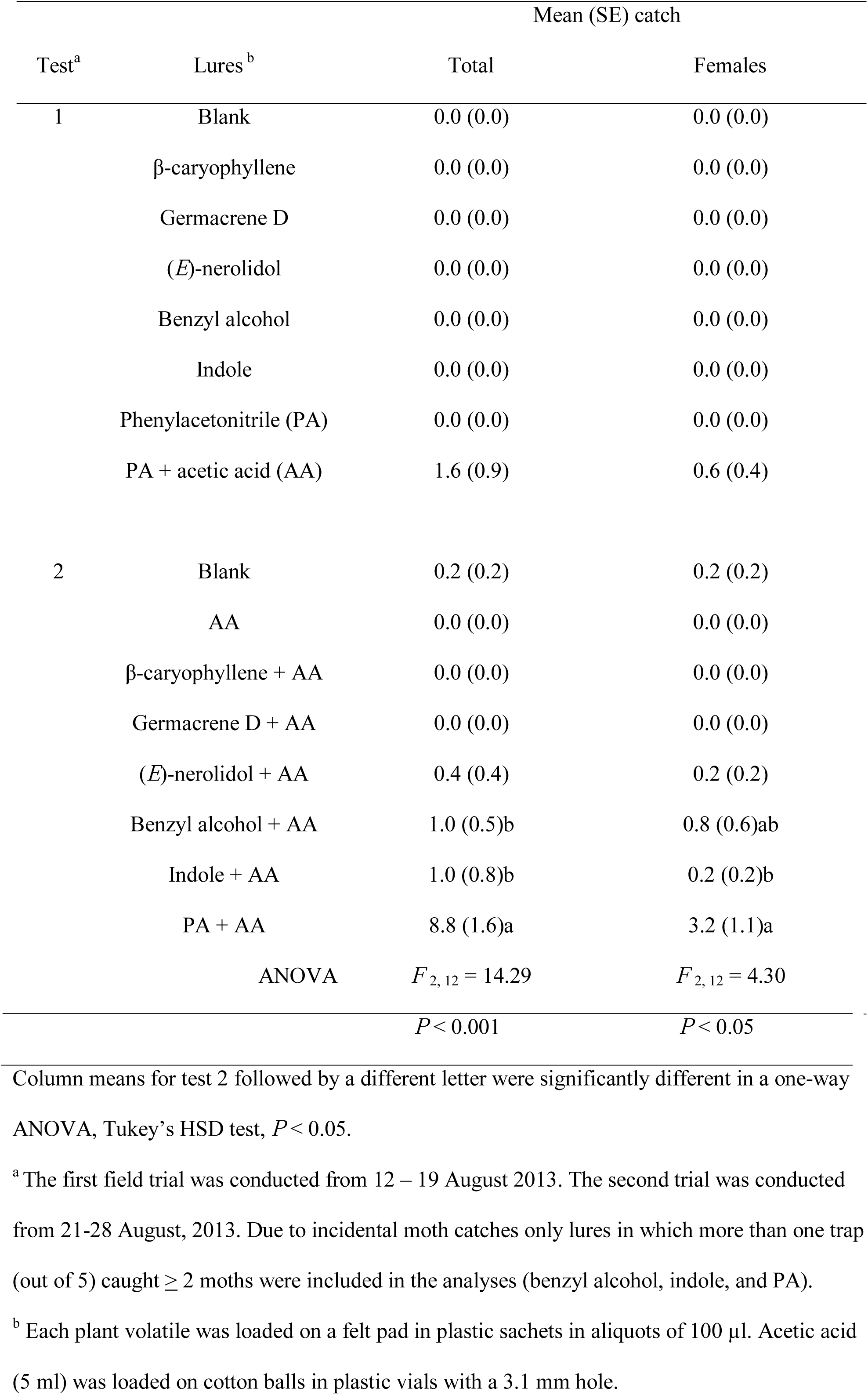
Comparison of catches of *Pandemis* leafroller in traps baited with different host-plant volatiles alone (test 1) or in combination with acetic acid (test 2), N = 5, Moxee, WA, 2013

### Combination lures with phenylacetonitrile

Consistent data were collected in both 2012 and 2013 when comparing catches of *C. pomonella* with phenylacetonitrile alone and with binary and ternary blends including pear ester and/or acetic acid (Tables 2 and 3). Pear ester in combination with acetic acid caught significantly more total and female *C. pomonella* than other lures. Adding phenylacetonitrile to pear ester and acetic acid significantly reduced both total and female moth catches of *C. pomonella.* Catches of *C. pomonella* with either phenylacetonitrile alone and in binary blends with either acetic acid or pear ester were low and did not differ significantly. In addition, traps with phenylacetonitrile alone in the 2013 test caught numbers of *C. pomonella* similar to numbers captured by either acetic acid or pear ester alone. Adding phenylacetonitrile to traps baited with the sex pheromone-pear ester combination lure (Pherocon CM-DA) plus acetic acid did not significantly reduce male, female, or total moth catches in 2014 (Fig. 1).

**Fig. 1.**
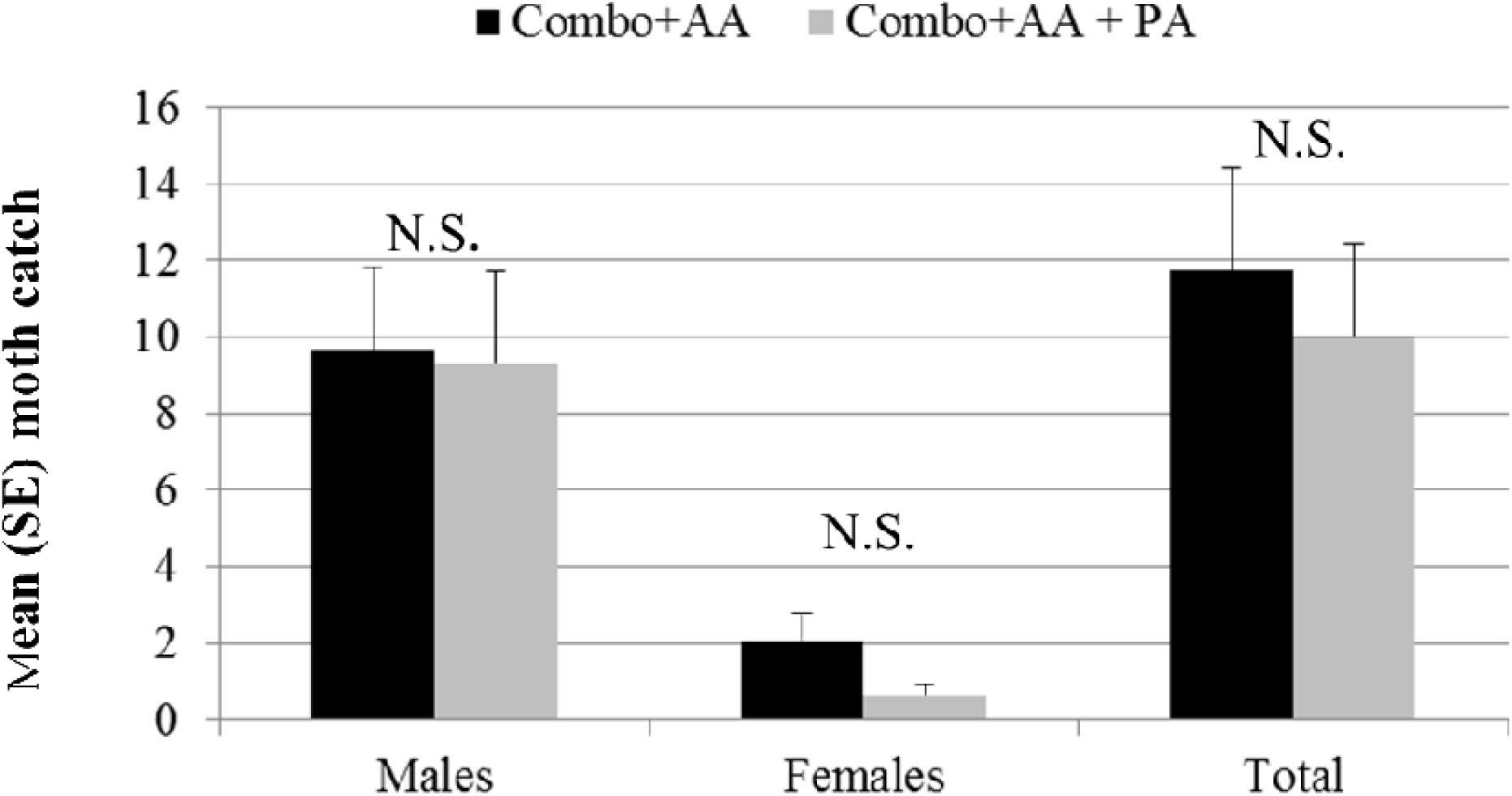
L Mean (SE) number of male, female, and total *C. pomonella* caught in traps with a sex pheromone + pear ester combo lure and an acetic acid co-lure versus these same lures with the addition of a phenylacetonitrile sachet lure, 7 – 21 July 2014, Moxee WA. ‘N.S.’ denotes a nonsignificant difference in moth catches between the two types of lures, *P* > 0.05.

**Table 2.**
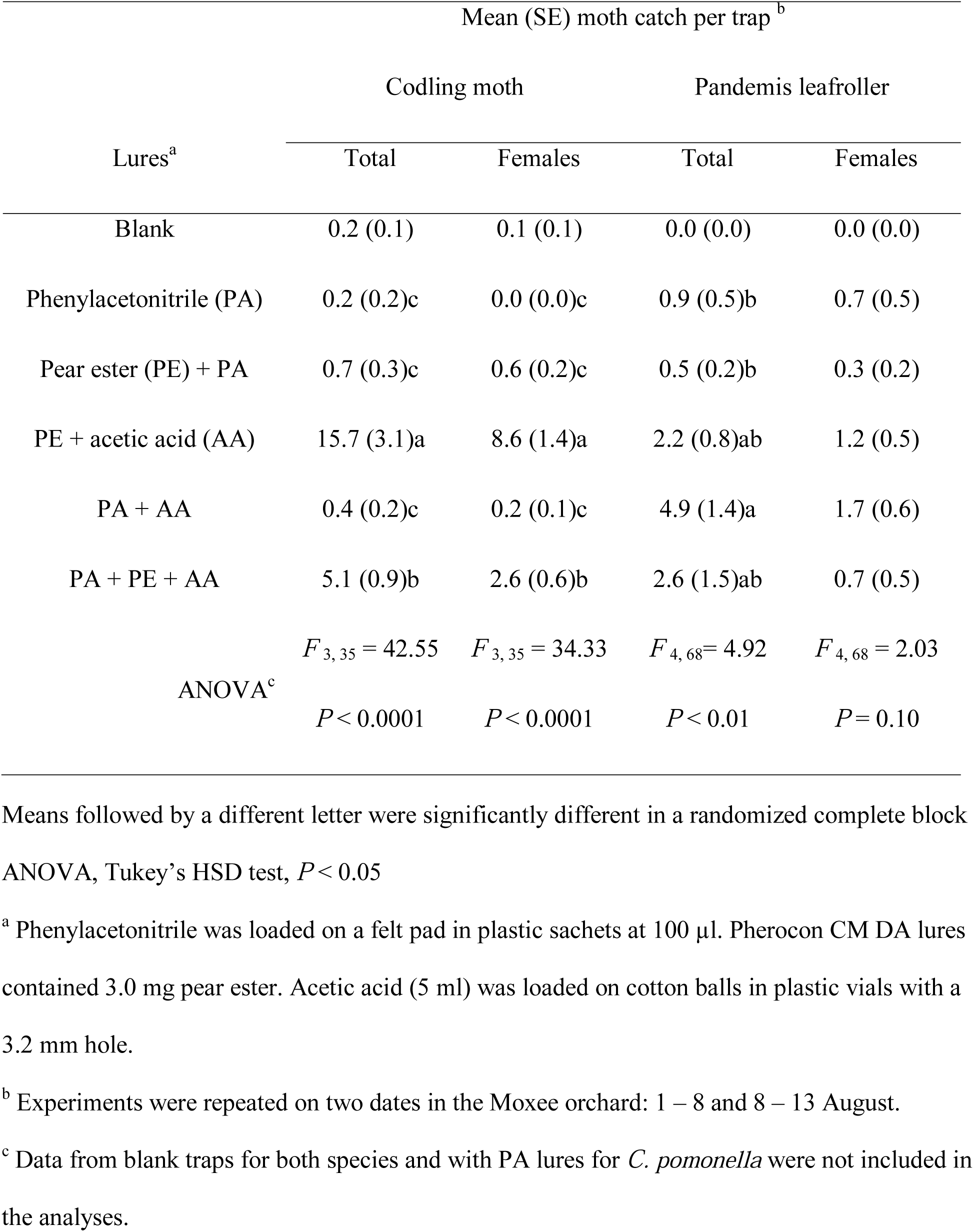
Comparison of catches of codling moth and *Pandemis* leafroller in orange delta traps unbaited or baited with combinations of phenylacetonitrile, pear ester, and acetic acid, N =10 for codling moth and N= 15 for Pandemis leafroller, Naches, WA 2012

**Table 3.**
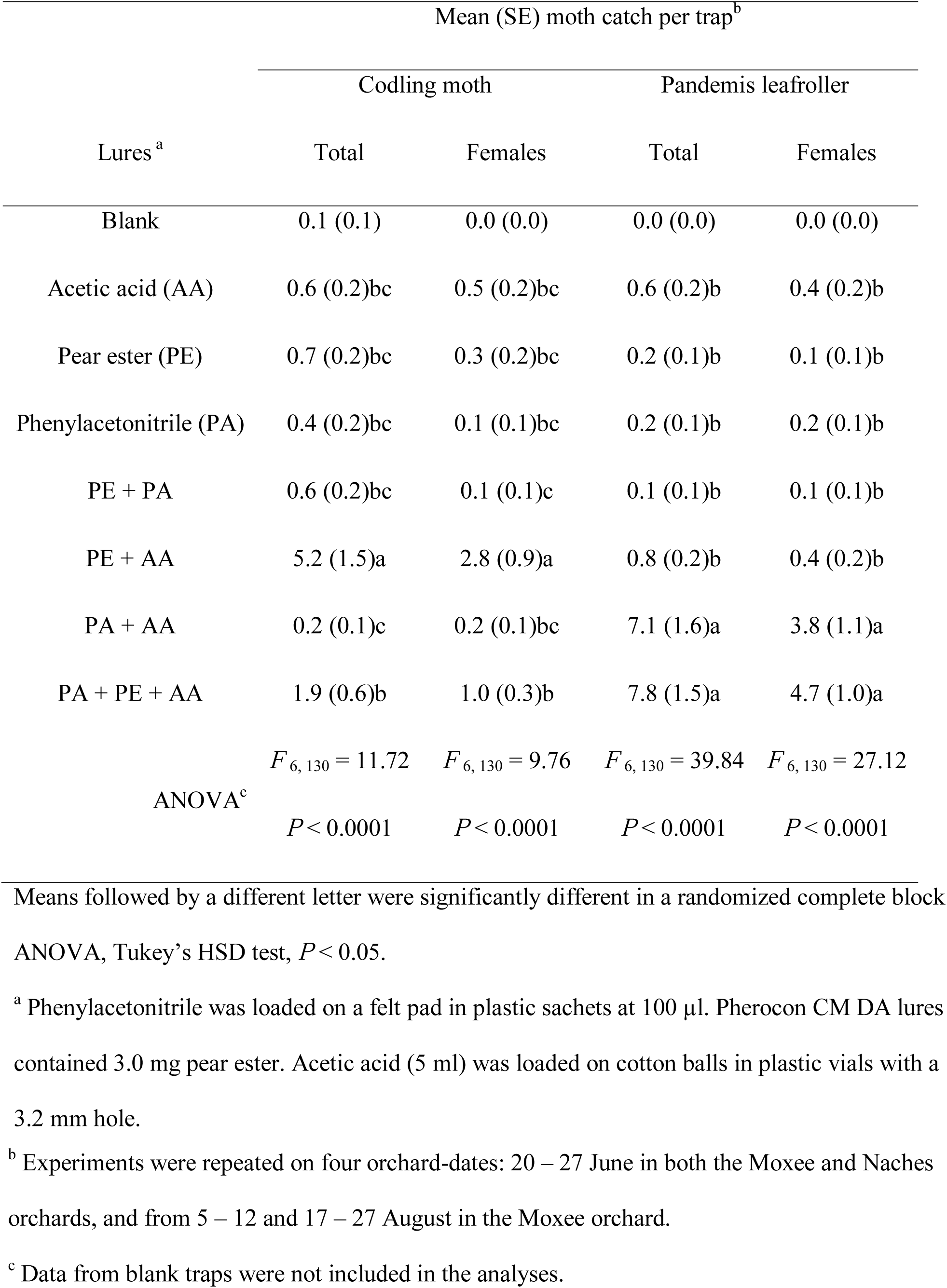
Comparison of catches of codling moth and *Pandemis* leafroller in orange delta traps unbaited or baited with combinations of phenylacetonitrile, pear ester, and acetic acid, N = 20, Moxee, WA 2013

Counts of *P. pyrusana* were more variable in traps than captures of codling moth between both years of the study (Tables 2 and 3). During 2013, traps baited with phenylacetonitrile plus acetic acid or in combination with pear ester caught significantly more total and female moths than all other treatments and did not differ among themselves. However, moth catches in 2012 were lower than in 2013 and no significant differences among treatments were found for female moth catch. Also, total moth catch in traps baited with pear ester plus acetic acid in 2012 was not significantly different from the four treatments that included phenylacetonitrile.

The average evaporation rate of acetic acid from the co-lures used with phenylacetonitrile was a significant factor affecting moth catches of *P. pyrusana* (Table 4). Evaporation rates ranged 18-fold across the three dispensers tested. Moth catches were significantly lower in traps with acetic acid lures with the lowest evaporation rate (plastic cup lure), and did not differ between traps baited with plastic vials with the two different hole sizes. Sachet lures loaded with phenylacetonitrile evaporated an average (SE) of 6.3 (0.2) mg d^−1^.

**Table 4.** Comparison of catches of *Pandemis* leafroller in orange delta traps baited with phenylacetonitrile plus one of three different acetic acid co-lures, N = 15, Moxee WA, 2013.

| Type of acetic acid lure<sup>a</sup> | Mean (SE) daily weight loss (mg) from lure | Mean (SE) weekly moth catch<sup>b</sup> |  |
| --- | --- | --- | --- |
|  |  | Total | Females |
| Plastic cup | 4.9 (0.1)c | 10.2 (2.2)b | 3.3 (0.8)b |
| 1.0-mm vial | 16.1 (2.0)b | 26.8 (7.2)a | 9.1 (2.4)a |
| 3.2-mm vial | 88.6 (3.3)a | 17.1 (3.5)a | 6.8 (1.4)a |
| ANOVA | $F_{2,27} = 500.8$<br>$P < 0.0001$ | $F_{2,40} = 12.56$<br>$P < 0.001$ | $F_{2,40} = 10.42$<br>$P < 0.001$ |

Means followed by a different letter were significantly different in a randomized complete block ANOVA, Tukey's HSD test, $P < 0.05$ .
<sup>a</sup> Acetic acid lures included the proprietary Pherocon AA (Trécé Inc., Adair, OK) plastic cup and polypropylene vials filled with 5 ml of acetic acid and with either 1.0 or 3.2 mm holes.
Phenylacetoneitrile sachet lures released an average (SE) of 6.3 (0.2) mg d<sup>-1</sup>.
<sup>b</sup> Experiments were repeated on three dates: 28 August - 4 September, 4 - 11 September and 12 - 19 September, 2013 in the Moxee orchard.

Moth catches of *P. pyrusana* in traps placed in the Naches orchard were much higher during 2014 than in the two previous years (Fig. 2). Total moth catch was significantly different and nearly 6-fold higher in traps baited with phenylacetonitrile than with the sex pheromone of *P. pyrusana (F*_1_, _8_ = 54.64, *P* < 0.001). Mean catch of both male (*F*_1_, _8_ = 9.30, *P* < 0.05) and female (*F* _1_, _8_ = 20.01, *P* < 0.01) moths alone were both significantly higher with this binary lure than the catch of males in the sex pheromone-baited traps.

**Fig. 2.**
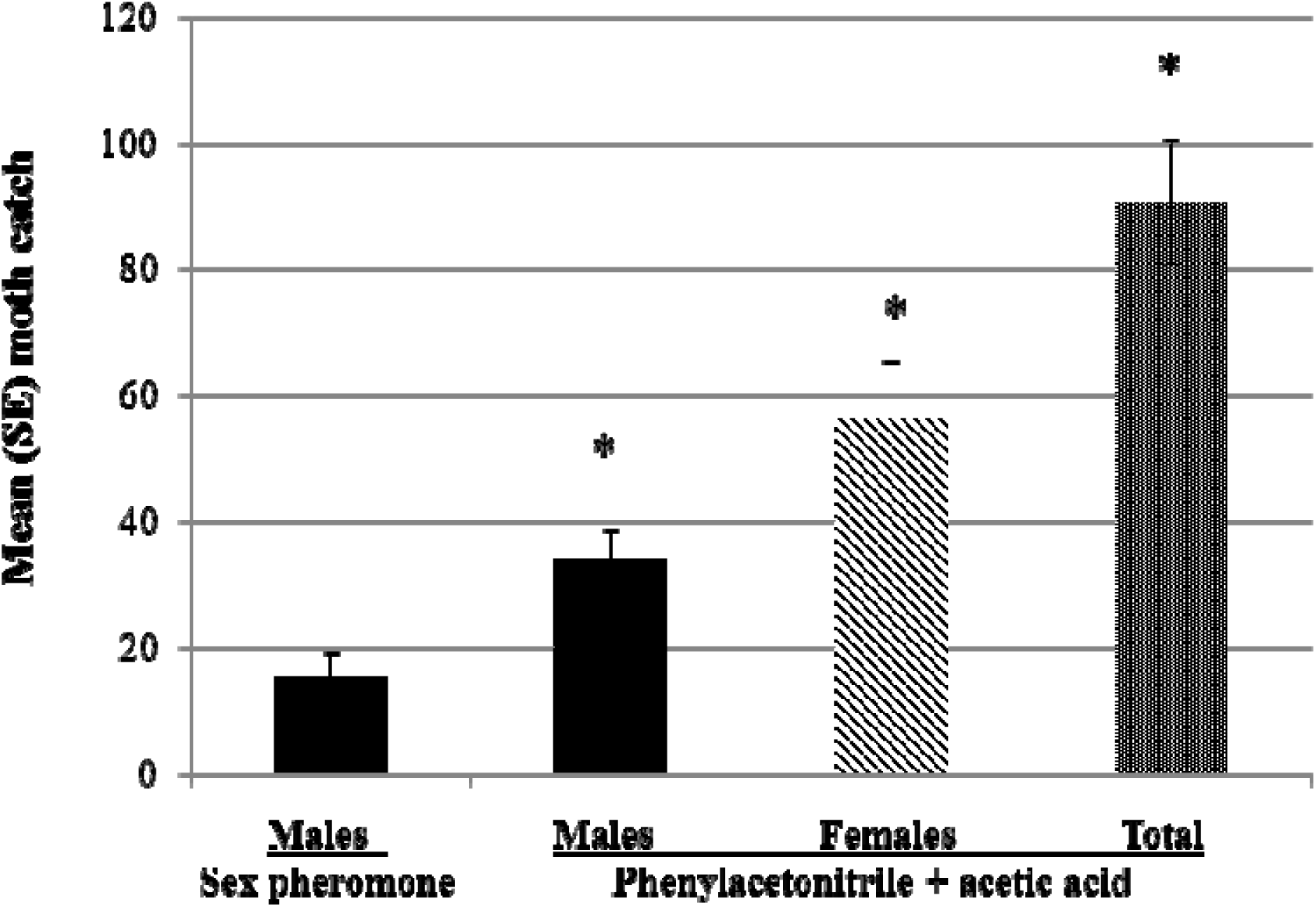
Mean (SE) number of male, female, and total *P. pyrusana* caught in traps baited with either the sex pheromone lure for *P. pyrusana* (males only) or with phenylacetonitrile plus an acetic acid co-lure (males, females, and total moths). ‘*’ denotes a significant mean difference with male catch in the sex pheromone-baited trap, *P* < 0.05.

## Discussion

In the studies reported here, phenylacetonitrile and other plant volatiles known to be induced by insect feeding, including benzyl alcohol, indole, and acetic acid provided no intrinsic attraction of *P. pyrusana* when tested alone. However, when any of the three compounds combined with acetic acid all, and especially phenylacetonitrile, exhibited strong attractiveness to adult male and female *P. pyrusana.* Benzyl nitrile and acetic acid has been reported as herbivore induced plant volatile in apple as results of LBAM larval feeding, while benzyl nitrile was identified as herbivore induced plant volatile in apple as results of OBLT and ESBM larval feeding (El-Sayed et al., 2016). In our previous work conducted in apple orchards in Canada,combination of benzyl nitrile and acetic acid or 2-phenylethanol and acetic acid were attractive to adult male and female OBLR and ESBM. The results obtained in this study combined with our earlier results with other tortricids suggest the positive response of adult tortricid to herbivores to HIPV compounds indicate that this is widespread phenomena among these species. Phenylacetonitrile has previously been studied with several tortricids and host-plant material in laboratory studies. For example, phenylacetonitrile was one of 22 compounds identified from the headspace (< 0.5%) of excised peach shoots, *Prunus persica* L. (Natale et al. 2003), but was not found to be attractive to mated female *Grapholita molesta* (Busck), when used in a blend with three green leaf volatiles plus benzaldehyde (Piñero and Dorn 2007). Apple leaves also released phenylacetonitrile within a blend of volatiles in response to larval feeding by the tortricids *E. postvittana* (Suckling et al. 2012) and *P. heparana* (Giacomuzzi et al. 2013). Yet, neither study measured whether it was attractive to these herbivorous insects.

Phenylacetonitrile is a relatively well known semiochemical involved in a range of insect-insect and insect-plant interactions. These include serving as a male-produced anti-aphrodisiac in the butterfly *Pieris brassicae* L.; facilitating a phoretic dispersal and arrestment by female egg parasitoid *Trichogramma brassicae* Bezdenko (Fatouros et al. 2005, Huigens et al. 2009); and acting as a male-specific repellant in the locust, *Schistocerca gregaria* (Forskal) in order to manage sperm competition within insect aggregations (Seidelmann and Ferenz 2002).

Phenylacetonitrile is also released by male and female flowers of boxelder trees, *Acer negundo* L., and it alone attracts nymphs, and both sexes of the adult box elder bug, *Boisea rubrolineata* (Barber) (Schwarz et al. 2009). Phenylacetonitrile was isolated from oilseed rape, *Brassica napus L*., and shown to attract cabbage seed weevil, *Ceutorhynchus assimilis* Payk (Smart and Blight 1997). Similarly, phenylacetonitrile is induced by feeding of the Japanese beetle, *Popillia japonica* Newman, and the fall webworm, *Hyphantria cunea* Drury, on several plant species, including crabapple, *Malus sp.*, and within a blend of induced volatiles, serves as an adult aggregation kairomone (Loughrin et al. 1995). Phenylacetonitrile was shown to be released from cabbage, *Brassica* spp., and nasturtium, *Tropaeolum majus* L., following larval feeding of *P. brassicae* and *P. rapae* L., but not from undamaged plants (Geervliet et al. 1997). In apple, a significant difference in the release of phenylacetonitrile was found among four cultivars, including, two transgenic scab resistant cultivars (Vogler et al. 2010).

Previous studies with plant-feeding Lepidoptera (primarily noctuids and pyralids) have correlated the presence of larval feeding on a host plant with the repulsion of the con-specific adults (Landolt 1993, De Moraes et al. 2001, Signoretti et al. 2012, Reisenman et al. 2013, Zakir et al. 2013, Hatano et al. 2015). No previous studies on tortricid moths have reported whether adults cue to host-plant volatiles induced by larval feeding. When presented alone the induced plant volatiles examined in this study exhibited no attraction to adult *P. pyrusana* based on the absence of moths in traps. However, we did not study whether these induced volatiles were behaviorally repellent or interfered with volatile-mediated host selection and/or sex pheromone communication (Hatano et al. 2015).

Our findings suggest that acetic acid is an important synergist of herbivore-induced plant volatiles for adult *P. pyrusana.* Interestingly, acetic acid has rarely been reported in the analysis of volatile collections from damaged apple foliage: trace levels were found in samples of cut flowering apple branches (Bengtsson et al. 2001), from apple leaves infested with phytophagous mites (Llusia and Peñuelas 2001), and from drought-stressed trees (Ebel et al. 1995).

Interestingly, an earlier paper found phenylacetonitrile was released by spider mite damage on apples (Takabayashi et al. 1994),. Acetic acid is a primary metabolite of plant metabolism under stress conditions and is regulated by the production and catabolism of ethanol and acetaldehyde (Oikawa and Lerdau 2013). However, acetic acid could also be produced by the growth of yeast and bacteria prevalent in the phyllosphere of apple leaves, or on surfaces of senescent flowers which are relatively rich in sugars and other carbohydrates (Atlas and Bartha 1981, Kinkel 1997). A potential role for microorganisms in the herbivore induction of plant volatiles has been suggested previously (Dicke and Hilker 2003, Hilker and Meiners 2006); yet, few studies have tried to measure this interaction. One exception was the purported role of regurgitated endosymbionts in the induction of plant defenses (Spiteller et al. 2000).

The interplay of microbial and plant volatiles with an adult tortricid pest has perhaps best been studied with *Lobesia botrana* (Denis and Schiffermüller) on grapes, *Vitis vinifera* L., (Tasin et al. 2005, 2011). In this system, neither acetic acid nor the induced volatile 2-phenylethanol was collected from undamaged flowers and fruit (Tasin et al. 2005). However, both compounds were abundant in volatiles from fruit infested with yeasts or acetic bacteria, and both female oviposition and net reproductive rate were significantly increased as a result of microbial infestations of fruit (Tasin et al. 2011). In contrast, fruit infected by *Botrytis cinerea* (Persoon), did not release acetic acid and was repulsive to females and reduced the overall fitness of the insect. Inoculation of fruit with acetic bacteria produced > 100-fold higher levels of acetic acid than with the yeasts, and oviposition was reduced compared with clean fruit, suggesting an overdose of acetic acid can occur. In our studies, the higher emission rates of acetic acid tested were more attractive, but neither the emission rates of phenylacetonitrile nor acetic acid have been optimized for attraction of *P. pyrusana*.

Our studies can likely impact the pest management of *P. pyrusana*. Surprisingly, traps baited with phenylacetonitrile and acetic acid caught more moths than sex pheromone traps in an orchard not treated with leafroller sex pheromones. These data may suggest that the commercial lures for *P. pyrusana* are not optimized or that other species in the *Pandemis limitata* (Robinson), group with different sex pheromone blends are more dominant in the two orchards studied (Dombroskie and Sperling 2012). The respective sex pheromones of *P. limitata* and *P. pyrusana* have been identified as 91:9 and 94:6 blends of (Z)-11-tetradecenyl acetate and (*Z*)-9-tetradecenyl acetate (Roelofs et al. 1976, 1977). At least one commercial lure has been reported to be relatively ineffective for *P. pyrusana* due to contamination with < 1.0% of (Z)-9-dodecenyl acetate (Brunner and Fisher 1998).

We found that adding pear ester to traps baited with phenylacetonitrile and acetic acid did not reduce the catch of *P. pyrusana.* However, adding phenylacetonitrile to traps baited with pear ester and acetic acid significantly reduced catches of *C. pomonella.* Thus ternary blends in a single trap would be useful in monitoring *P. pyrusana,* but may compromise catches of *C. pomonella.* However, a significant drop in catch of *C. pomonella* did not occur when phenylacetonitrile and acetic acid were added to the combination sex pheromone plus pear ester lure. The potential benefit of improved monitoring of *P. pyrusana,* especially of female moths, with phenylacetonitrile would have to outweigh the information lost from a drop in catches of female *C. pomonella* in these combination traps (summarized in Knight et al. 2014). Hypothetically, this could occur in certified-organic orchards if the pest pressure from *C. pomonella* is low, efficacious tools to manage leafrollers are limited, and especially, when sex pheromone-based mating disruption for leafrollers is used and the standard sex pheromone-baited traps are ineffective. The ability to monitor multiple species could fuel the adoption of labor-saving remote electronic traps (Guarnieri et al. 2011, Kim et al. 2011).

Our positive results with enhancing the bisexual moth catches of *P. pyrusana* suggest that studies should continue to identify and test other plant volatiles induced by insect feeding on pome fruits (Boevé et al. 1996, Suckling et al. 2012, Giacomuzzi et al. 2013). The use of more complex and ratio-specific blends of induced volatiles should be evaluated, perhaps both alone and in the context of the common volatiles released by uninfested and infested plants (Bruce and Pickett 2011). For example, green leaf volatiles have been found to enhance catches with sex pheromones in field trials with some tortricid pests (Light et al. 1993, Varela et al. 2011, Yu et al. 2015). The release rates of both (Z)-3-hexenyl acetate and (Z)-3-hexenyl benzoate were increased with herbivory (Suckling et al. 2012), and in a previous study, combining (*Z*)-3-hexenyl acetate with acetic acid nearly doubled the catch *of P. pyrusana* (Knight et al. 2014). Polyphagous tortricid pests such as *P. pyrusana* attack a number of horticultural crops other than pome fruits, and future studies should more broadly characterize the induced plant volatiles in these crops to identify other possible attractants, including green leaf volatiles, aromatics, alcohols, and terpenes, and test these in combination with acetic acid as attractive lures.

## Acknowledgements

We would like to thank Duane Larson and Chey Temple, Agricultural Research Service, Wapato, WA for their technical assistance in the laboratory and field. We would like to acknowledge our appreciation to Bill Lingren, Trécé Inc., Adair, OK for donating the commercial lures used in our studies. This project was supported with partial funding from the Washington Tree Fruit Research Commission, Wenatchee, WA.

